# Looking for mimicry in a snake assemblage using deep learning

**DOI:** 10.1101/789206

**Authors:** Thomas de Solan, Julien Pierre Renoult, Philippe Geniez, Patrice David, Pierre-Andre Crochet

## Abstract

Batesian mimicry, with its highly colorful species and astonishing mimic-model resemblance, is a canonical example of evolution by natural selection. However, Batesian mimicry could also occur in inconspicuous species and rely on subtle resemblance. Although potentially widespread, such instances have been rarely investigated, such that the real frequency of Batesian mimicry has remained largely unknown. To fill this gap, we developed a new approach using deep learning to quantify the resemblance between putative mimics and models from photographs. We applied this method to quantify the frequency of Batesian mimicry in Western Palearctic snakes. Potential mimics were revealed by an excess of resemblance with sympatric venomous snakes compared to random expectations. We found that 8% of the non-venomous species were potential mimics, among which all were imperfect mimics. This study is the first to quantify the frequency of Batesian mimicry in a whole community of vertebrates, and shows that even concealed species can be reliably identified as potential models. Our approach should prove useful to detect mimicry in other communities, and more generally it highlights the benefits of deep learning for quantitative studies of phenotypic resemblance.

## 1. Introduction

Batesian mimicry occurs when palatable or harmless species resemble an unprofitable (e.g. unpalatable or dangerous) model species, adopting some of its characteristics, and consequently gaining protection from predators (Bates 1862). Mimicry is a canonical example of evolution through natural selection; it was already discussed by Darwin with Wallace (Darwin 1887) and still constitutes a proliferous field of research in ecology and evolution (Sherratt 2002; Nishikawa et al. 2015; Joshi et al. 2017; Bosque et al. 2018). Nevertheless, some fundamental aspects of this defensive strategy remain poorly known, notably its frequency in plants and animals, mainly because detecting mimicry is no simple matter.

Historically, research on Batesian mimicry has long been confined to a few spectacular cases, like hoverflies (Penney et al. 2012), butterflies (Kitamura and Imafuku 2015) or kingsnakes (Brodie and Janzen 1995), which possess conspicuous colors and show a resemblance between models and mimics so striking that it cannot be fortuitous. This perfection itself is a double-edged sword. It served as an evidence of the Darwinian evolution, but it also provided a distorted picture of what mimicry can be. Indeed, neither conspicuousness nor high resemblance is a necessary condition for Batesian mimicry. Actually, even when their overall appearance is inconspicuous or camouflaged, venomous species may eventually be spotted by predators, which can recognize species using cues that have not necessarily evolved as specific signals. Harmless and palatable species can mimic these cues to gain additional protection from mimicry while being themselves cryptic (Barlow and Wiens 1977; Mound et al. 2005; Gianoli and Carrasco-Urra 2014; Corcobado et al. 2016). Moreover, experimental evidence suggest that an imperfect resemblance between a mimic and its model, known as imperfect mimicry, can still provide protection against predators (Dittrich et al. 1993). Thus, most examples found in literature do not reflect the potential diversity of mimicry in nature. Focusing on these classic cases is likely to bias our perception of mimicry, making us underestimate its frequency.

If we want to fully understand the role of mimicry in shaping phenotypic evolution in natural communities, we need to go beyond the few exceptionally perfect cases to expose common and imperfect products of natural selection. The detection of mimicry within a community requires a quantification of the resemblance between species. In mimicry studies, two main categories of methods have been traditionally used to estimate resemblance. The first methods analyze predefined features; for example morphometric measures, the geometry of color patterns or the distribution of orientations and spatial frequencies in light changes in images (Penney et al. 2012; Taylor et al. 2016; Troscianko et al. 2017; Endler et al. 2018). These methods usually attempt to model physiological perceptual processes and thus can provide information on the role of these processes in the evolution of mimicry. However, by being confined to the studied processes, they may be limited for capturing the high dimensionality of the feature space relevant for assessing visual resemblance (Zhang et al. 2018). The second category of methods relies on psychological tests with human subjects (e.g., Cuthill and Bennett 1993). These methods can provide more accurate assessments of overall perceptual resemblance; however, extensive tests and very large sample size are necessary to mitigate the individual variations between evaluators (Ishihara et al. 2001; Kitayama et al. 2003).

Artificial Intelligences, and especially deep convolutional neural networks (hereafter DNNs), offer new opportunities to model perceptual resemblance (Jozwik et al. 2017; Cichy and Kaiser 2019). For example, a psychophysical experiment conducted with thousands of humans showed that DNN’s measure of resemblance between complex stimuli (e.g., pictures of a crowd) predict the human averaged perceived resemblance better than any previous models (Zhang et al. 2018). Moreover, DNNs can estimate resemblance despite noisy variations, that is, between stimuli that differ in position, orientation, occlusion and illumination (Jozwik et al. 2017). Thus, DNNs are highly suited to exploit the nearly inexhaustible resource of non-standardized photographs posted on the World Wide Web. DNNs have revolutionized many scientific fields in recent years, including biology (e.g. in genomics: Zhou and Troyanskaya 2015, medicine: Shen et al. 2017) but the potential of DNN as a model of human and animal perception, which can handle very large databases of non-standardized data, has been overlooked in visual ecology thus far.

Several cases of Batesian mimicry between venomous and non-venomous snakes of the Western Palearctic (Europe, North Africa and the Middle-East) has been reported in literature, sometime based on experimental validation or mimic behavior but mainly using a human subjective assessment of the visual resemblance between potential model and mimic (Gans 1961; Werner and Frankenberg 1982; Werner 1983; Valkonen et al. 2011*a*; Aubret and Mangin 2014; Valkonen and Mappes 2014). None of the venomous model species in the study area possesses highly conspicuous colors, and most of them are cryptic in their environment (Turk 1958; Heatwole and Davison 1976; Werner 1983). However the zigzag dorsal patterns of vipers has been shown to elicit avoidance in bird predators (Valkonen et al. 2011*b*) in addition to providing camouflage (Santos et al. 2014). In spite of these case studies, no attempt has been made to evaluate the extent of mimicry in the Western Palearctic snake assemblage. Western Palearctic snakes thus appear as a good model to measure the frequency of mimicry in situations more complicated than most classic examples described in the literature. Here we incorporated DNN into a workflow to identify potential mimics among all 122 snake species known in the Western Palearctic at the time of the study.

To identify candidate mimics, we developed a method that quantifies the visual resemblance between candidate mimics (in our example, the non-venomous Western Palearctic snakes) and candidate models (venomous Western Palearctic snakes). We (i) trained DNNs to recognize the candidate (venomous) models before (ii) using the DNNs to assess the resemblance between candidate mimic (non-venomous) and candidate models. To identify candidate mimics, we (iii) combined the information on resemblance with information on distribution to detect non-venomous species that exhibit more phenotypic resemblance to one or several models that live in sympatry than expected by chance. We found that 8% of the non-venomous species were candidate mimics, among which almost all were imperfect mimics to a human eye. A global test of the hypothesis that mimicry affected the evolution of phenotype in the Western Palearctic snake assemblage was highly significant. We discuss this result in the light of the known ecology of Western Palearctic snakes and of the validity of modeling non-human predator vision using DNNs and regular images.

## 2. Materials and Methods

We analyzed all snake species of the Western Palearctic (after the taxonomic list of Geniez 2015), 122 in total, with 35 venomous species and 87 non-venomous species. Table S1 lists all species of the Western Palearctic indicating taxonomy, venomousness, the number of images analyzed and the source of distribution data.

### (1) Image datasets

We created two image datasets, one for the non-venomous snakes (838 images; mean = 9.6 images per species) and one for the venomous snakes (1802 images; mean = 50.0 per species; details in Table S1), with more images than average for the venomous species exhibiting high intraspecific polymorphism. Because images are rare for several of the studied species, our datasets may appear small compared to those of other studies using DNNs. However, recent investigations revealed that, with an appropriate training setup (see *transfer learning* below), DNNs can reach very high performances even when trained on limited dataset (Körschens et al. 2018) –this is confirmed by our results (see below).

Some snake species consist of several well-separated morphotypes. As DNNs can solve nonlinear problems, we did not separate morphotypes. Rather, we expected the models to learn that morphotypes represent a single species. We included images of distinct individuals except for a few species for which only a few images were available. In addition to the 35 species of Western Palearctic venomous snakes (each representing a distinct class of pictures), we added one class of “foreign” venomous snakes to the venomous dataset. This class was made of 80 photographs of 80 different species living outside the Western Palearctic but in climates similar to those of the study area, i.e. temperate and desert (list of the species used in Table S1). We selected only images showing a clear dorsal view of the whole specimen (see Fig. 2 for examples). Eighty percent of the images were collected on the Internet, the remaining being taken in the field by the authors or colleagues. An expert herpetologist (PG) validated all identifications individually.

**Figure 1.**
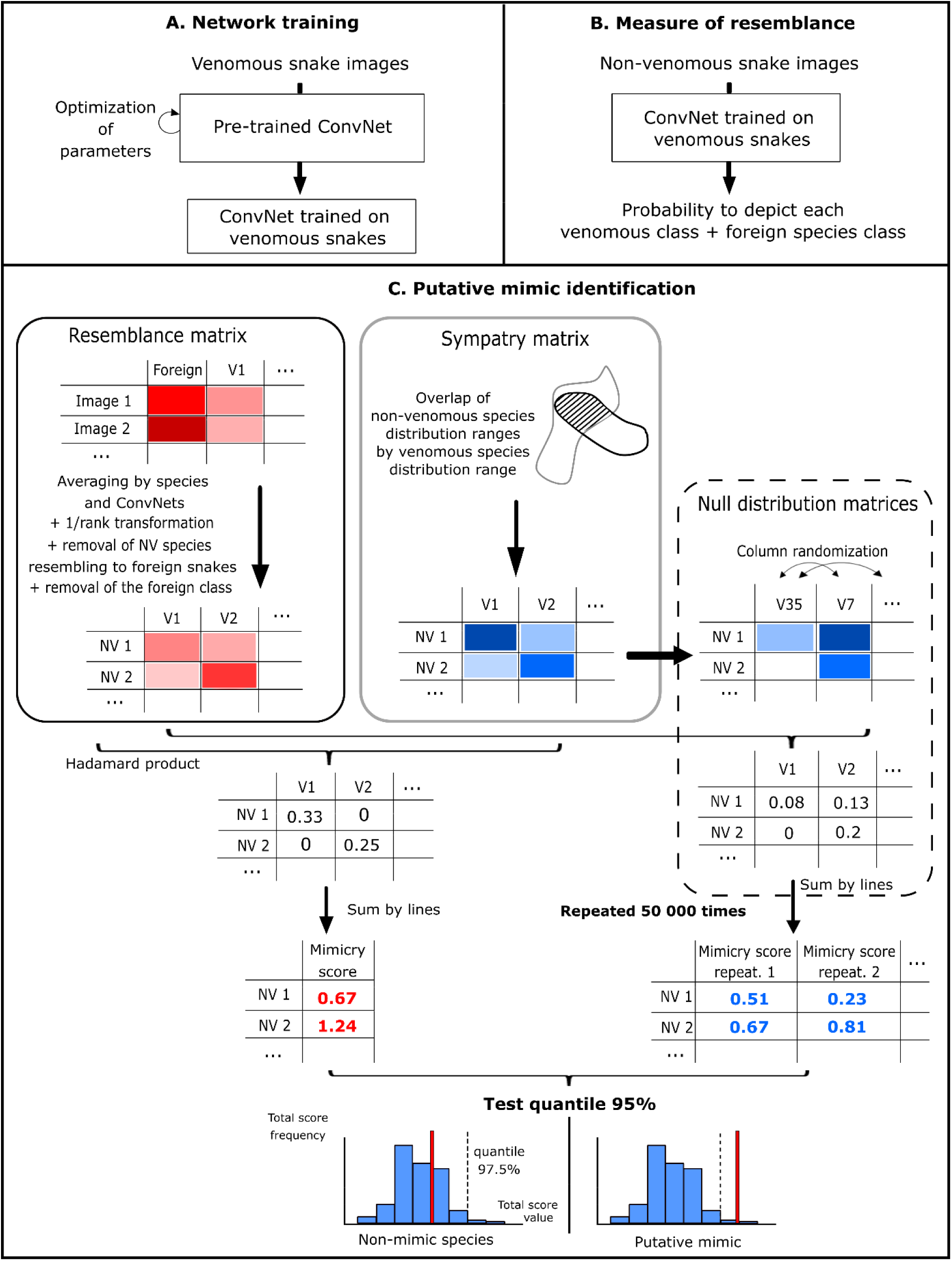
Workflow of the three-step method used to detect candidate mimic species among non-venomous snakes and candidate model species among venomous ones. (A) Training of the artificial neural networks to recognize 35 venomous snake species plus the “foreign” venomous snake class. (B) Measuring the resemblance of non-venomous snakes (one measure per image) with each species of venomous snakes plus the “foreign” class. (C) Building the resemblance matrix and the sympatry matrix, and testing an excess of resemblance between sympatric species using a null distribution to identify candidate mimics.

**Figure 2.**
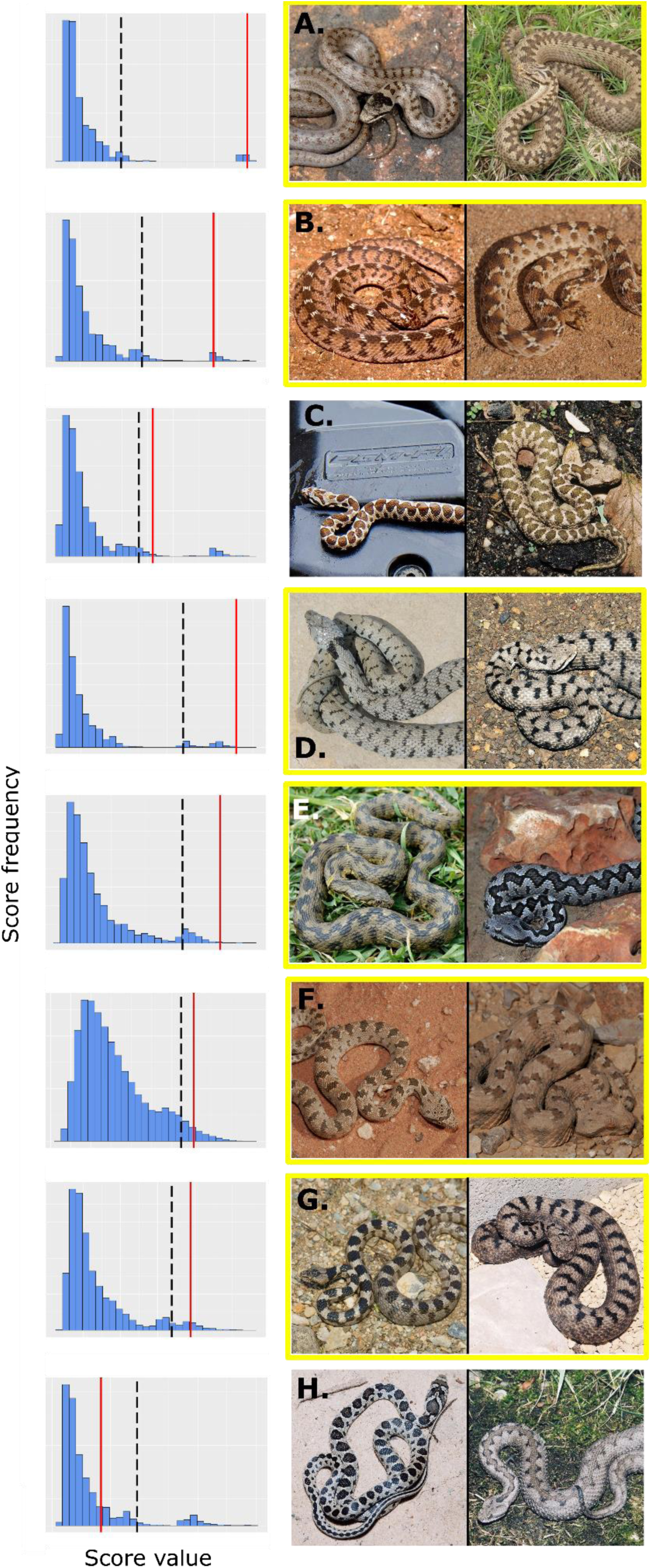
The seven mimic and model species identified in our analysis (A-G) and one example pair of sympatric but non-resembling species (H), with mimicry scores (red), quantiles 95% (black dashed line) and the null distribution (blue) of each species pairs. Species framed in yellow have been previously reported as mimetic in the literature (Gans 1961; Werner and Frankenberg 1982; Werner 1983; Valkonen et al. 2011*a*; Aubret and Mangin 2014). (A) *Coronella austriaca – Vipera berus*. (B) *Dasypeltis sahelensis – Echis pyramidum*. (C) *Hemorrhois nummifer – Montivipera xanthina*. (D) *Natrix helvetica – V. aspis*. (E) *N. maura – V. latastei*. (F) *Spalerosophis diadema – Pseudocerastes fieldi*. (G) *Telescopus fallax – V. ammodytes.* (H) *H. hippocrepis – V. latastei.* Photo credits: Matthieu Berroneau, Olivier Buisson, Luis García Cardenete, Alexandre Cluchier, Philippe Geniez, André Langenbach, Jean Muratet, Gilles Pottier and Jean-François Trape.

### (2) Geographic distribution dataset

We constructed distribution ranges from three types of data: (1) when available, we used distribution maps downloaded from the IUCN Red List website (www.iucnredlist.org, accessed on February 2018); (2) otherwise, we reconstructed by hand the spatialized distribution maps from pictures of distribution ranges found on the http://apps.who.int/bloodproducts/snakeantivenoms/database/ website and from other specialized literature; (3) when no distribution map was available, we built maps from occurrence data found in the specialized literature (references listed in Table S1). We used QGIS 2.18 (Quantum GIS Development Team 2013) to create the distribution maps. Each distribution map was depicted by a spatialized polygon projected in WGS 84.

### (3) Methodological workflow

DNNs were first trained to recognize the different venomous species of the Western Palearctic plus the “foreign” class (Figure 1A). Trained DNNs were then asked to classify pictures of non-venomous snakes as venomous species plus the “foreign” category (Figure 1B). DNNs provided, for each image, a probability of being attributed to each venomous species class. The “foreign” class avoids forcing the networks to attribute an image to one of the potential model even if it does not resemble any of the Western Palearctic venomous species. For each non-venomous species, attribution probabilities were then averaged over all available pictures to build a matrix of resemblance between non-venomous and venomous species with averaged attribution probabilities as entries. These probabilities were transformed into resemblance scores, i.e. in inversed ranks of probability (1/rank, in ascending order; Figure 1C). Thus, the classes of venomous snakes to which one particular nonvenomous snake was attributed most often, i.e. ranking first or second, had the highest resemblance score. This step allows to reduce the prediction variability that can arise between DNNs due to differences of architectures or training algorithms (Sameen and Pradhan 2017; Kim et al. 2019). Next, we deleted from this matrix all non-venomous species for which the “foreign” class received the highest resemblance score, thus discarding species that were clearly not mimics, as well as the foreign species class. In parallel, we built a sympatry matrix of the same size as the resemblance matrix, which gives the proportion of the geographic distribution of each non-venomous species that overlaps with the distribution of each venomous species. Following a previous study (Miller et al. 2019), we considered that the proportion of sympatry reflected the potential extent of selection for mimicry. This, in fact, is an approximation as for a few allopatric species pairs, distribution ranges may be close enough to allow predators associating both snake species. Yet most allopatric pairs are distant by hundreds or thousands of kilometers in our dataset, which limits this possibility (Pfennig and Mullen 2010). The last step of our workflow was to calculate “total scores” by summing, for each non-venomous species, the resemblance scores weighted (Hadamard product) by sympatry scores.

Our method did not include phylogenetic correction for the measure of the resemblance because all the venomous and non-venomous snakes belong to two distant and monophyletic clades. The resemblance between these clades thus cannot be attributed to phylogenetic relationship.

Candidate mimics and models were identified statistically by comparing total scores to a null distribution. This null distribution was generated by randomly shuffling the venomous species in the sympatry matrix before calculating the resemblance score for each non-venomous species, and by replicating this operation 50,000 times (Figure 1C). Species for which the total scores fell in the right tail (e.g., the 5% highest scores) of the null distribution were more similar to sympatric venomous species than expected by chance and were thus considered candidate mimics (and their sympatric venomous species considered candidate models).

We tested the same hypothesis of an excess of resemblance multiple times; thus, we tested whether significant results could have arisen by chance. We used the Fisher’s method, also named Fisher’s combined probability test (Fisher 1925), to test if there was, across all the non-venomous species, a significant excess of resemblance with the sympatric venomous snakes. We combined the *p*-values of the 27 non-venomous species that were not classified as foreign venomous snakes into a *X*^2^ statistic and compared it to a chi-squared distribution with 2 × 27 degrees of freedom. This effectively provides a global test of the hypothesis that mimicry affected the evolution of phenotype in the Western Palearctic snake assemblage. All statistical analyses were performed with R (Development Core Team 2008).

### (4) Image pre-processing

Snake images were squared (by cropping around snakes) and downsized to 299 × 299 pixels. The venomous snake dataset was split into two: 90% of images were used for training and 10% for validation (i.e., to monitor performances while tuning training hyperparameters). We used an image augmentation procedure to improve invariance to noisy variations and to limit overfitting: before every training epoch, images were rotated with an angle sampled between 0° and 180°, randomly flipped horizontally and vertically, up to 20% zoomed in or out, and RGB color channel randomly shifted by 0.2. Images of non-venomous snakes were squared and downsized only.

### (5) DNN training and predictions

Training a DNN to recognize snake species from scratch would have required a very large dataset of images to achieve good performances. Following other studies (e.g., Norouzzadeh et al. 2018), we rather used “transfer learning”, in which a DNN pre-trained on a very large dataset is fine-tuned with a new, smaller dataset. In this study, we transferred the Xception network (Chollet 2017) pre-trained on ImageNet (a dataset of 14 million images belonging to more than 20 000 classes; Deng et al. 2009). We tested other networks also trained on ImageNet (ResNet, Alexnet), which all yielded qualitatively similar results but reached lower performance in species recognition. To limit overfitting, we replaced the original final classification layer by a first fully-connected convolutional layer (2048 units), a dropout layer (dropout probability: 0.7), a second fully-connected convolutional layer (256 units), a second dropout layer (dropout probability: 0.7) and a 36-ways softmax classifier. The highest performances were reached by fine-tuning all layers (with a very small learning rate) rather than the last, added layers only. We trained three networks separately using three different optimizers: Nadam (learning rate 0.00003, decay 0), Single Gradient Descent (learning rate 0.001, decay 0.00001, Nesterov momentum 0.9) and Adagrad (learning rate: 0.0005, decay: 0) (Zheng 2009; Ju et al. 2017). The training was stopped when we visually estimated that the curves of validation loss did not decrease further for 20 consecutive epochs, i.e. after 50 epochs for training with Stochastic Gradient Descent, 120 epochs with Adagrad and 45 epochs with Nadam (Figure S3).

We estimated the classification probabilities of each non-venomous snake image with each of the three DNNs trained with different optimizers, before averaging probabilities across DNNs. The whole training and prediction procedure was repeated 15 times, to produce 15 matrices of resemblance scores that were eventually averaged into a single resemblance matrix. All DNN analyses were performed with the Keras library (Chollet 2015) in Python language.

### (6) Test of background influence

To confirm that network predictions were effectively based on the snakes, and not on the background, we replaced the background in 60 images. Snakes were cut out and pasted onto photos of ground only. Ground photos depicted rock, dead leaves, plants and sand. Of the 60 snake images, 20 belonged to venomous snakes, 20 to non-venomous candidate mimic snakes and 20 to non-venomous non-candidate mimic snakes.

We used the 3×15 trained networks to predict the resemblance scores for the 60 images with a replaced background. We used non-parametric Multivariate Analysis Of Variance, with adonis function of vegan package (Dixon 2003) in R, to test the effect of background replacement on resemblance scores. We controlled variation between species by entering species names as an explanatory variable. We checked for multivariate homoscedasticity using the betadisper function of vegan package (non-significant, effect of background: F = 0.15, p = 0.69).

## 3. Results

After training, DNNs were able to recognize the species of a Western Palearctic venomous snake with an accuracy of 77% ± 0.004 SE (evaluated on the validation dataset; random guess: 2.78% accuracy) and a top-3 accuracy of 92% ± 0.015 SE (probability that one of the three classes with the highest prediction matches the correct class), indicating that DNNs learned the most distinctive traits of each species. This is a remarkably high performance considering the limited size of the training dataset, and the extreme resemblance among several vipers, which makes their identification challenging even for experts when the geographic origin is unknown (Geniez 2015). In comparison, a previous DNN-based analysis of species recognition based on 35,200 training images from the Internet representing 289 species of reptiles reached 45.9% and 80.9% top-1 and top-5 accuracy, respectively (Van Horn et al. 2018).

Comparing weighted resemblance scores to the null distribution, with a 5% threshold, our method identified seven non-venomous species out of the 87 (8%) as candidate mimics of venomous snakes: these species have a higher resemblance with their sympatric venomous species than with a random assemblage of venomous snakes (list of potential mimics and models, Figure 2). Fisher’s method confirmed the overall significance of excessive resemblance with sympatric venomous snakes across the tested non-venomous species set (*X*^*2*^ = 109.3, p < 0.001). In addition, resemblance scores were not driven by similarity between image backgrounds, as they were not affected by shuffling backgrounds among images (non-parametric MANOVA performed with 60 images, background effect: F = 0.74, p = 0.67).

## 4. Discussion

We propose here a new automatic method to infer potential mimetic pairs and to evaluate the frequency of mimicry within an entire group of organisms based on non-standardized photographs. We applied it to quantify for the first time the extent of mimicry in an assemblage of vertebrates at a continental scale (see Wilson et al. 2015 for a related work with invertebrates). Although the hypothesis of mimicry would have to be validated experimentally for each particular species pair, our analysis confirms the existence in Western Palearctic snakes of a pattern typical of Batesian mimicry (i.e. pairwise resemblance between sympatric pairs of unrelated nontoxic and toxic preys), and suggests that this defensive strategy has influenced their phenotypic evolution. Finally, our analysis confirms previous researches in demonstrating that DNNs can reach high level of accuracy even when trained on small datasets (Hansen et al. 2018; Körschens et al. 2018; Mathis et al. 2018).

Several arguments from the natural history of snakes support the validity of our method to identify mimics and models. Firstly, six of the seven species identified as potential mimics by our method have been mentioned as such in the literature (Gans 1961; Werner and Frankenberg 1982; Werner 1983; Valkonen et al. 2011*a*; Aubret and Mangin 2014; see Figure 2). Secondly, the mimicry hypothesis is supported for all the detected species by additional evidence of mimicry, based on behavioral or/and acoustic traits. The putative mimics *Natrix helvetica, N. maura*, C*oronella austriaca, Spalerosophis diadema* and *Telescopus fallax* displace their maxillary bones to produce a triangular head shape when disturbed, increasing visual resemblance with their viper model (Werner and Frankenberg 1982; Werner 1983; Valkonen et al. 2011*a*; Aubret and Mangin 2014). In addition, *Natrix maura* also produces a hissing sound similar to the sympatric *Vipera aspis* and *V. berus* (Aubret and Mangin 2014) while the non-venomous *Dasypeltis sahelensis* and its sympatric venomous *Echis pyramidum* possess the same modified scales on the flanks that are rubbed to produce a similar warning signal when disturbed (Gans 1961). Finally, even if it has never been reported yet as a potential mimic in the literature, *Hemorrhois nummifer* shows the same behavioral and acoustic warning signal as its sympatric vipers, *Montivipera xanthina* and *Macrovipra lebetina* (authors’ personal observation).

Remarkably, we found that even well camouflaged species (e.g., vipers of the genus *Pseudocerastes*; Figure 2) could be a model for non-venomous species. The lack of any pattern or color signaling venomousness in this species confirms that mimicry can be effective without conspicuous traits, and advocate for a better separation between the concepts of Batesian mimicry, communication signaling and conspicuousness. The link between these three concepts seems to represent a historical legacy of the easier detection of mimicry when it involves conspicuous aposematic models. The method presented here, which uses DNN-based resemblance coupled with information on geographic co-occurrence, avoids a biased selection favoring conspicuous organisms when studying Batesian mimicry.

An additional advantage of our DNN-based approach is its ability to suggest mimicry even when resemblance is imperfect. Most of the species identified by our method are imperfect mimics, at least from our human point of view, as their colors and patterns match only partially those of their model (Figure 2). Although any of the eleven non-exclusive hypotheses proposed in the literature to explain imperfect mimicry (see Kikuchi and Pfennig 2013; Dalziell and Welbergen 2016 for a detailed list) could apply to Western Palearctic snakes, only the “back-up signal” hypothesis is currently supported by ecological evidence. This hypothesis suggests that imperfect mimicry can be reinforced by complementary signals (Johnstone 1996). It is convincing especially for mimics that show similarities in behavioral, morphological and acoustic warning signals with their sympatric venomous species (Gans 1961; Werner 1983; Valkonen et al. 2011*a*; Aubret and Mangin 2014). In addition to the eleven hypotheses, one characteristic of the studied snakes could help explain how imperfect mimicry can evolve. Some model species are highly variable, which may prevent predators from associating a specific phenotype to venomousness (Kikuchi and Pfennig 2010) and thus make them more careful and less likely to attack snakes with ambiguous phenotypes. This explanation is plausible at least for the three candidate models *Vipera ammodytes, V. beru*s and *V. aspis*, which exhibit significant intraspecific and often intra-population variation in coloration (Figure S2). More research is needed to evaluate the validity of these different explanations and to determine why imperfect mimicry seems ubiquitous among Western Palearctic snakes.

Although additional cues comfort the mimicry hypothesis for all detected species, our method can only be used for suggesting candidate mimics and models, and a proper demonstration of mimicry would require additional data. For instance, mimetic species are likely to have similar size and ecology than their models. This information was not provided by our photographs but may influence the plausibility of the mimicry hypothesis for some species (see below). Moreover, most studies of interspecific resemblance, including our method, are fundamentally limited in their ability to distinguish between mimicry and other factors of resemblance. For example, phylogenetic relatedness is a classical cause of resemblance; however this hypothesis does not apply to our study as venomous and non-venomous snakes belong to two distant, reciprocally monophyletic clades. Resemblance could also arise by ecological convergence because of similar abiotic environmental constraints (Rajpurohit et al. 2008), or similar selective pressures for camouflage (Stevens et al. 2013). These factors could lead to overestimating the frequency of mimicry in our results. This issue potentially affects all mimicry studies that do not test predator behavior in a natural environment. However, because most studies focus on conspicuously colored and presumably aposematic species (Brodie and Janzen 1995; Cheney and Marshall 2009; Penney et al. 2012) researchers usually assume that resemblance can only be attributed to mimicry. This assumption is seldom tested although even conspicuous colors may have camouflage function (Mochida et al. 2015; Barnett et al. 2018). In Western Palearctic snakes, body colors are not conspicuous. It would thus be interesting to perform predation tests in natural conditions to investigate whether the features copied by potential mimics are those effectively used by predators to recognize models (e.g., as in Kristiansen et al. 2018), and thus to confirm that resemblance is effectively driven by mimicry. Nevertheless, as discussed previously all identified mimics use behavioral or acoustic imitation in addition to color pattern and, as resemblance in several ontogenetically independent traits can hardly be attributed to ecological convergence, to our view mimicry remains the most likely explanation.

The ability to detect imperfect and non-conspicuous mimicry makes our method less conservative compared to methods entirely based on human detection, for evaluating the frequency of mimicry. Even so, we may have failed to identify potential mimic-model pairs. First, the 5^th^ percentile threshold in the null distribution used to interpret resemblance scores is entirely conventional and may overlook some mimicry systems. We found two species below but close to the 5^th^ percentile threshold. The first species, *Dolichophis jugularis*, displays the same black coloration as its sympatric species *Walterinnesia morgani*, but it has a different timing of activity (one is diurnal while the other is nocturnal) and is thus probably not a mimic. The second species, *Rhagerhis moilensis*, resembles the sympatric *Cerastes cerastes* in coloration but has a different size and shape (Geniez 2015). Mimicry hypothesis is implausible, but cannot be discounted for this species. Therefore, the 5^th^ percentile threshold should be interpreted cautiously and we recommend not drawing firm conclusions without examining species close to a decision threshold. Second, we may have failed to identify candidate mimics and models because, for the non-venomous polymorphic species (e.g. Leviton 1970; Kark et al. 1997; Mebert 2011), we could not test mimicry for each morphotype separately. Instead, we had to analyze all the morphotypes together, and test mimicry at the species level. This complicates the detection of mimicry as only some morphotypes of these polymorphic species may mimic venomous snakes (e.g., in *Hemorrhois ravergieri*, Werner and Frankenberg 1982, see below). Testing mimicry for each morphotype is possible knowing the precise geographical distribution of each morphotype, but this information was not available for most polymorphic species. These two limits may lead to an under-estimation of the mimicry frequency in natural communities.

Among the species suggested as mimetic in literature, three were not identified in our study (Gans 1961; Werner and Frankenberg 1982; Werner 1983). One of them, *Hemorrhois ravergieri* is considered a mimic of the sympatric *Vipera ammodytes transcaucasiana* (Werner and Frankenberg 1982). Our method failed to identify it as such probably because it exhibits a strong polymorphism, with only some of its variants thought to mimic vipers (Werner and Frankenberg 1982). A second species, *Dasypeltis bazi*, could have been missed by our approach because of the limited number of pictures in our dataset (six in total, this rare and poorly-known species was recently described by Saleh and Sarhan 2016) and the marked variation in coloration between individuals. However, the third unrecognized mimic, *Telescopus dhara* (Werner 1983), is not polymorphic, and we believe that claims for mimicry in this species are too optimistic, as it bears no particular resemblance with the proposed model species, *Echis coloratus*, even from visual inspection.

Our study highlights the benefits of DNNs for studying resemblance between phenotypes in ecology and evolution. DNNs have been previously used in ecology to analyze big data, e.g., to identify species from camera trap images (Norouzzadeh et al. 2018). However, their capacity to model biological perception has been thus far overlooked (Cichy and Kaiser 2019). A growing body of literature in humans has shown that DNNs predict the perceived resemblance between complex stimuli better than all other models, even those specifically tuned with neurophysiological data (Yamins et al. 2014; Kriegeskorte 2015; Kubilius et al. 2016). Of particular relevance for the study of camouflage and mimicry, one study showed that whichever perceptual task a DNN is trained for (e.g., recognizing objects from images, segmenting foreground in video frames), DNN accurately predicts perceptual similarity even near the discrimination threshold (Zhang et al. 2018).

In mimicry systems (and more generally in most communication systems) the relevant observer is not human. Are the performances of DNNs at reproducing human perception also relevant to other animals? These performances are based on three characteristics (Kriegeskorte 2015). The first one, the capacity of DNNs to solve high-level tasks (Zhang et al. 2018), is shared by all animals. The second is their general architecture (a stack of convolution, activation and pooling functions) which reproduces operations performed sequentially and iteratively by neurons in the different human visual areas (Güçlü and van Gerven 2015). Similar operations are also found, beyond humans, in at least all Vertebrates (Renoult et al. 2019). The third is the structure of input data. In this study, we used RGB images that match human trichromatic vision (ie based on three cone types). Other animals have different cone types (e.g., two in carnivores, four in birds), tuned to different wavelengths. How these differences affect DNN models of perceptual space is a nearly unexplored avenue of research (but see Bergeron and Fuller 2018). Yet indirect evidence suggests that the impact would be relatively minor. Indeed, studies on bird plumages that used both RGB images and bird-specific cone stimulations showed no qualitative differences (Dale et al. 2015; Miller et al. 2019). Accordingly, several authors have argued that RGB images provide major insight into ecology, evolution, and behavior in nonhuman species (Seddon et al. 2010; Bergeron and Fuller 2018). In addition, in our particular case, the use of ranks to express relative resemblance makes our mimicry-detection method insensitive to how precisely the similarity measure produced by DNNs scales to similarity perceived by animals.

A potentially important limitation of RGB images, however, is to eliminate wavebands to which humans –but not other vertebrates– are blind. The ultraviolets (UV), in particular, may be problematic. Regarding our study, the main predators of snakes in the Western Palearctic are non-primate mammals and raptors (Valkonen et al. 2011*b*): they have a very limited UV vision and they most certainly do not use UV cues for foraging (Jacobs 2009; Lind et al. 2013, 2014). Thus, UV-based mimicry seems unlikely in snakes. More generally, however, one should keep in mind that DNNs will never detect resemblance from information not present in the data.

Last, in our study we analyzed non-standardized images. Although standardization removes irrelevant information from the data, we think that non-standardized images are ecologically more relevant. Indeed, predators experience substantial variation in perceptual conditions, in relation to snake position, background, shading, light intensity and color. This variation could affect the evolution of mimic phenotypes; for example, it has been suggested to explain imperfect mimicry (relaxed selection hypothesis, Kikuchi and Pfennig 2013). While RGB pictures from the Internet may not precisely match variation experienced by predators, drastically limiting all variation (by standardization) would make the dataset un-natural. Standardized images would be nevertheless useful in other contexts, e.g. to determine which visual features contribute the most to deceive predators.

To conclude, Artificial Intelligence, here represented by DNNs, has a great potential for the study of mimicry. We used it to detect candidate Batesian mimics and models among all Western Palearctic snakes. We found that Batesian mimicry is a likely defensive strategy for 8% of non-venomous snakes. Although there is still room for methodological improvement, DNNs already allow to measure resemblance while accounting for the high dimensionality of visual phenotypes, simultaneously accounting for colors, texture, patterns and proportions, and without requiring standardized data that may be difficult to collect for many species. Through their ability to detect imperfect and non-conspicuous mimicry, DNNs further avoids the bias of many mimicry studies towards colorful and spectacularly resembling species. Finally, beyond mimicry, the ability to handle complex visual patterns makes DNNs highly promising tools for modeling resemblance in general, with broad perspectives of applications in evolution and ecology.

