## Supplementary material for "Looking for mimicry in a snake assemblage using deep learning": table_S1_figures_S2_S3

**Table S1. Number of photographs, rational for species exclusion and source of distribution maps for each of the 35 venomous species and 87 non-venomous species of the Western Palearctic snakes, plus the 80 foreign species of the foreign class.**

| Venomousness | Species name | Number of images | Rational for species exclusion | Distribution map sources |
|---|---|---|---|---|
| | *Atractaspis engaddensis* | 41 | | (1) |
| | *Bitis arietans* | 40 | | (2) |
| | *Cerastes cerastes* | 53 | | (3) |
| | *Cerastes gasperettii* | 40 | | (1) |
| | *Cerastes vipera* | 50 | | (1) |
| | *Daboia mauritanica* | 48 | | (1) |
| | *Daboia palaestinae* | 54 | | (1) |
| | *Echis carinatus* | 45 | | (3) |
| | *Echis coloratus* | 40 | | (3) |
| | *Echis pyramidum* | 52 | | (1) |
| | *Gloydius halys* | 40 | | (3) |
| | *Macrovipera lebetina* | 45 | | (3) |
| | *Montivipera bornmuelleri* | 40 | | (1) |
| | *Montivipera bulgardaghica* | 81 | | (1) |
| | *Montivipera raddei* | 40 | | (1) |
| | *Montivipera wagneri* | 40 | | (1) |
| | *Montivipera xanthina* | 47 | | (1) |
| | *Naja haje* | 40 | | (3) |
| | *Naja nubiae* | 41 | | (3) |
| | *Pseudocerates fieldi* | 40 | | (1) |
| | *Vipera ammodytes* | 84 | | (1) |
| **Venomous** | *Vipera anatolica* | 45 | | (1) |
| | *Vipera aspis* | 84 | | (1) |
| | *Vipera berus* | 64 | | (3) |
| | *Vipera darevskii* | 40 | | (1) |
| | *Vipera dinniki* | 41 | | (1) |
| | *Vipera eriwanensis* | 40 | | (1) |
| | *Vipera graeca* | 40 | | (4) |
| | *Vipera kaznakovi* | 58 | | (1) |
| | *Vipera latastei* | 81 | | (1) |
| | *Vipera renardi* | 52 | | (5) |
| | *Vipera seoanei* | 70 | | (1) |
| | *Vipera ursinii* | 40 | | (1) |
| | *Walterinnesia aegyptia* | 40 | | (1) |
| | *Walterinnesia morgani* | 26 | | (3) |
| | **Foreign species class :** | | | |
| | *Acanthophis cryptamydros* | | | |
| | *Acanthophis laevis* | | | |
| | *Acanthophis pyrrhus* | | | |
| | *Acanthophis wellsei* | | | |
| | *Antairoserpens warro* | | | |
| | *Antaresia childreni* | | | |
| | *Antaresia maculosa* | 80 | | |
| | *Antaresia perthensis* | | | |
| | *Austrelaps ramsayi* | | | |
| | *Austrelaps superbus* | | | |

| | |
|---|---|
| | *Boiga irregularis* |
| | *Brachyurophis australis* |
| | *Brachyurophis fasciolatus* |
| | *Brachyurophis incinctus* |
| | *Brachyurophis morrisi* |
| | *Brachyurophis roperi* |
| | *Brachyurophis semifasciatus* |
| | *Cacophis churchilli* |
| | *Cacophis krefftii* |
| | *Calamaria lautensis* |
| | *Cerberus rhynchops* |
| | *Crotalus adamanteus* |
| | *Crotalus catalinensis* |
| | *Crotalus mitchellii* |
| | *Crotalus oreganus* |
| | *Crotalus stejnegeri* |
| | *Crotalus tigris* |
| | *Cryptophis nigrescens* |
| | *Cryptophis nigrostriatus* |
| | *Cryptophis pallidiceps* |
| | *Demansia calodera* |
| | *Demansia olivacea* |
| | *Demansia psammophis* |
| | *Demansia reticulata* |
| | *Demansia rufescens* |
| | *Demansia torquata* |
| | *Demansia vestigiata* |
| **Venomous** | *Drysdalia mastersii* |
| | *Drysdalia coronoides* |
| | *Elapognathus minor* |
| | *Fordonia leucobalia* |
| | *Furina diadema* |
| | *Furina ornata* |
| | *Hemiaspis damelii* |
| | *Hoplocephalus bitorquatus* |
| | *Hoplocephalus stephensii* |
| | *Leiopython albertisii* |
| | *Liasis mackloti* |
| | *Liasis olivaceus* |
| | *Mixcoatlus browni* |
| | *Mixcoatlus melanurus* |
| | *Morelia oenpelliensis* |
| | *Notechis scutatus* |
| | *Oxyuranus microlepidotus* |
| | *Oxyuranus scutellatus* |
| | *Parasuta monachus* |
| | *Parasuta nigriceps* |
| | *Parasuta spectabilis* |
| | *Paroplocephalus atriceps* |
| | *Pseudechis australis* |
| | *Pseudechis colletti* |
| | *Pseudechis guttatus* |
| | *Pseudechis porphyriacus* |
| | *Pseudonaja affinis* |

| | | | |
|---|---|---|---|
| Venomous | *Pseudonaja aspidorhyncha* | | |
| | *Pseudonaja guttata* | | |
| | *Pseudonaja ingrami* | | |
| | *Pseudonaja modesta* | | |
| | *Pseudonaja textilis* | | |
| | *Pseudonaja mengdeni* | | |
| | *Rhinoplocephalus bicolor* | | |
| | *Simoselaps bertholdi* | | |
| | *Simoselaps minimus* | | |
| | *Stegonotus parvus* | | |
| | *Suta ordensis* | | |
| | *Suta punctata* | | |
| | *Tropidechis carinatus* | | |
| | *Tropidonophis mairii* | | |
| | *Vermicella intermedia* | | |
| | *Vermicella snelli* | | |
| Non venomous | *Boaedon fuliginosus* | 10 | (6–8) |
| | *Coronella austriaca* | 10 | (1) |
| | *Coronella girondica* | 10 | (1) |
| | *Dasypeltis bazi* | 6 | (9) |
| | *Dasypeltis sahelensis* | 10 | (6, 7) |
| | *Dolichophis caspius* | 10 | (1) |
| | *Dolichophis jugularis* | 10 | (1) |
| | *Dolichophis schmidti* | 10 | (1) |
| | *Eirenis aurolineatus* | 10 | (1) |
| | *Eirenis barani* | 10 | (1) |
| | *Eirenis collaris* | 10 | (1) |
| | *Eirenis coronella* | 10 | (1) |
| | *Eirenis coronelloides* | 10 | (1) |
| | *Eirenis decemlineatus* | 10 | (1) |
| | *Eirenis eiselti* | 10 | (1) |
| | *Eirenis levantinus* | 10 | (1) |
| | *Eirenis lineomaculatus* | 10 | (1) |
| | *Eirenis modestus* | 10 | (1) |
| | *Eirenis occidentalis* | 6 | (10) |
| | *Eirenis punctatolineatus* | 10 | (1) |
| | *Eirenis rothi* | 10 | (1) |
| | *Eirenis thospitis* | 10 | (1) |
| | *Elaphe dione* | 10 | (1) |
| | *Elaphe quatuorlineata* | 10 | (1) |
| | *Elaphe sauromates* | 10 | (1) |
| | *Eryx colubrinus* | 10 | (6, 7) |
| | *Eryx jaculus* | 10 | (6, 10) |
| | *Eryx jayakari* | 10 | (1) |
| | *Eryx miliaris* | 10 | (6, 10) |
| | *Hemorrhois algirus* | 10 | (1) |
| | *Hemorrhois hippocrepis* | 10 | (1) |
| | *Hemorrhois nummifer* | 10 | (1) |
| | *Hemorrhois ravergieri* | 10 | (1) |
| | *Hierophis cypriensis* | 10 | (1) |
| | *Hierophis gemonensis* | 10 | (1) |
| | *Hierophis viridiflavus* | 10 | (1) |
| | *Indotyphlops braminus* | Introduced | |

| | | | |
|---|---|---|---|
| Non venomous | *Letheobia episcopus* | 10 | (1) |
| | *Letheobia simonii* | 10 | (1) |
| | *Lytorhynchus diadema* | 10 | (1) |
| | *Macroprotodon abubakeri* | 10 | (1) |
| | *Macroprotodon brevis* | 10 | (1) |
| | *Macroprotodon cucullatus* | 10 | (1) |
| | *Macroprotodon mauritanicus* | 10 | (11) |
| | *Malpolon insignitus* | 10 | (12) |
| | *Malpolon monspessulanus* | 10 | (1) |
| | *Micrelaps muelleri* | 10 | (1) |
| | *Muhtarophis barani* | 10 | (6) |
| | *Myriopholis algeriensis* | 10 | (1) |
| | *Myriopholis cairi* | 3 | (6) |
| | *Myriopholis macrorhyncha* | 10 | (6) |
| | *Natrix astreptophora* | 10 | (13) |
| | *Natrix helvetica* | 10 | (14) |
| | *Natrix maura* | 10 | (14) |
| | *Natrix natrix* | 10 | (1) |
| | *Natrix tessellata* | 10 | (1) |
| | *Platyceps chesneii* | 7 | (15, 16) |
| | *Platyceps collaris* | 10 | (1) |
| | *Platyceps elegantissimus* | 10 | (1) |
| | *Platyceps florulentus* | 10 | (1) |
| | *Platyceps najadum* | 10 | (1) |
| | *Platyceps rhodorachis* | 10 | (6) |
| | *Platyceps rogersi* | 10 | (1) |
| | *Platyceps sinai* | 10 | (1) |
| | *Platyceps saharicus* | 10 | (6) |
| | *Psammophis aegyptius* | 10 | (17) |
| | *Psammophis schokari* | 10 | (17) |
| | *Psammophis sibilans* | 10 | (18) |
| | *Rhagerhis moilensis* | 10 | (6, 7) |
| | *Zamenis scalaris* | 10 | (1) |
| | *Rhynchocalamus dayanae* | 1 | (19) |
| | *Rhynchocalamus melanocephalus* | 10 | (1) |
| | *Rhynchocalamus satunini* | 10 | (19) |
| | *Spalerosophis diadema* | 10 | (6) |
| | *Spalerosophis dolichospilus* | 10 | (1) |
| | *Telescopus dhara* | 10 | (6, 20) |
| | *Telescopus fallax* | 10 | (6, 10, 20) |
| | *Telescopus hoogstrallii* | 10 | (1) |
| | *Telescopus nigriceps* | 10 | (1) |
| | *Telescopus obtusus* | 4 | (6, 20) |
| | *Telescopus tessellatus* | 8 | (1) |
| | *Telescopus tripolitanus* | 10 | (6, 20) |
| | *Xerotyphlops vermicularis* | 10 | (1) |
| | *Zamenis hohenackeri* | 10 | (1) |
| | *Zamenis lineatus* | 10 | (1) |
| | *Zamenis longissimus* | 10 | (1) |
| | *Zamenis persicus* | 10 | (1) |
| | *Zamenis situla* | 10 | (1) |

1. IUCN (2018) The IUCN Red List of Threatened Species. Version 2018-2. Available at: http://www.iucnredlist.org. Downloaded on 12 December 2018.
2. Barlow A, et al. (2013) Phylogeography of the widespread African puff adder (*Bitis arietans*) reveals multiple Pleistocene refugia in southern Africa. *Mol Ecol* 22(4):1134–1157.
3. World Health Organization (2010) Database of venomous snakes. Available at: http://apps.who.int/bloodproducts/snakeantivenoms/database/.
4. Mizsei E, et al. (2017) Nuclear markers support the mitochondrial phylogeny of *Vipera ursinii–renardi* complex (Squamata: Viperidae) and species status for the Greek meadow viper. *Zootaxa* 4227(1):75.
5. Ferchaud A-L, et al. (2012) Phylogeography of the *Vipera ursinii* complex (Viperidae): mitochondrial markers reveal an east-west disjunction in the Palaearctic region. *J Biogeogr* 39(10):1836–1847.
6. Geniez P (2018) *Snakes of Europe, North Africa and the Middle East : a photographic guide* (Princeton University Press).
7. Trape J-F, Mané Y (2006) Guide des serpents d'Afrique occidentale; savane et désert. 226.
8. Trape J-F, Mediannikov O (2016) Cinq serpents nouveaux du genre *Boaedon* Duméril, Bibron & Duméril, 1854 (Serpentes : Lamprophiidae) en Afrique centrale. *Bull la Société Herpétologique Fr* 159:61–111.
9. Saleh M, Sarhan M (2016) The egg-eating snake (Colubridae: *Dasypeltis*) of Faiyum, Egypt, with the description of a new species. *Bull la Société Herpétologique Fr* 160:25–48.
10. Safaei-Mahroo B, et al. (2015) The Herpetofauna of Iran: Checklist of Taxonomy, Distribution and Conservation Status. *Asian Herpetol Res* 6(4):257–290.
11. Wade E (2001) Review of the false smooth snake genus *Macroprotodon* (Serpentes, Colubridae) in Algeria with a description of a new species. *Bull Nat Hist Museum Zool Ser* 67:85–107.
12. Mangiacotti M, et al. (2014) Head shape variation in eastern and western Montpellier snakes. *Acta Herpetol* 9(2):167–177.
13. Kindler C, et al. (2018) Phylogeography of the Ibero-Maghrebian red-eyed grass snake (*Natrix astreptophora*). *Org Divers Evol* 18(1):143–150.
14. Kindler C, et al. (2017) Hybridization patterns in two contact zones of grass snakes reveal a new Central European snake species. *Sci Rep* 7(1):7378.
15. Schätti B, Tillack F, Kucharzewski C (2014) Platyceps rhodorachis (JAN, 1863) - A study of the racer genus *Platyceps* BLYTH, 1860 east of the tigris (Reptilia: Squamata: Colubridae). *Vertebr Zool* 64(3):297–405.
16. Schätti B, Schmitz A (2006) Re-assessing *Platyceps ventromaculatus* (Gray, 1834) (Reptilia: Squamata: Colubrinae). *Rev suisse Zool* 113:747–768.
17. Gonçalves DV, et al. (2018) The role of climatic cycles and trans-Saharan migration corridors in species diversification: Biogeography of *Psammophis schokari* group in North Africa. *Mol Phylogenet Evol* 118:64–74.

18. Kelly CMR, Barker NP, Villet MH, Broadley DG, Branch WR (2008) The snake family Psammophiidae (Reptilia: Serpentes): Phylogenetics and species delimitation in the African sand snakes (*Psammophis* Boie, 1825) and allied genera. *Mol Phylogenet Evol* 47(3):1045–1060.

19. Tamar K, Šmíd J, Göçmen B, Meiri S, Carranza S (2016) An integrative systematic revision and biogeography of *Rhynchocalamus* snakes (Reptilia, Colubridae) with a description of a new species from Israel. *PeerJ* 4:e2769.

20. Crochet P-A, et al. (2008) Systematic status and correct nomen of the western North African cat snake: *Telescopus tripolitanus* (Werner, 1909) (Serpentes: Colubridae), with comments on the other taxa in the dhara-obtusus group. *Zootaxa* 1703:25–46.

**Figure S2. Sample of the intraspecific variability observed in the same population of *Vipera aspis* present in the Briançonnais (France, province of Hautes-Alpes).** Photo credit: Grégory Deso

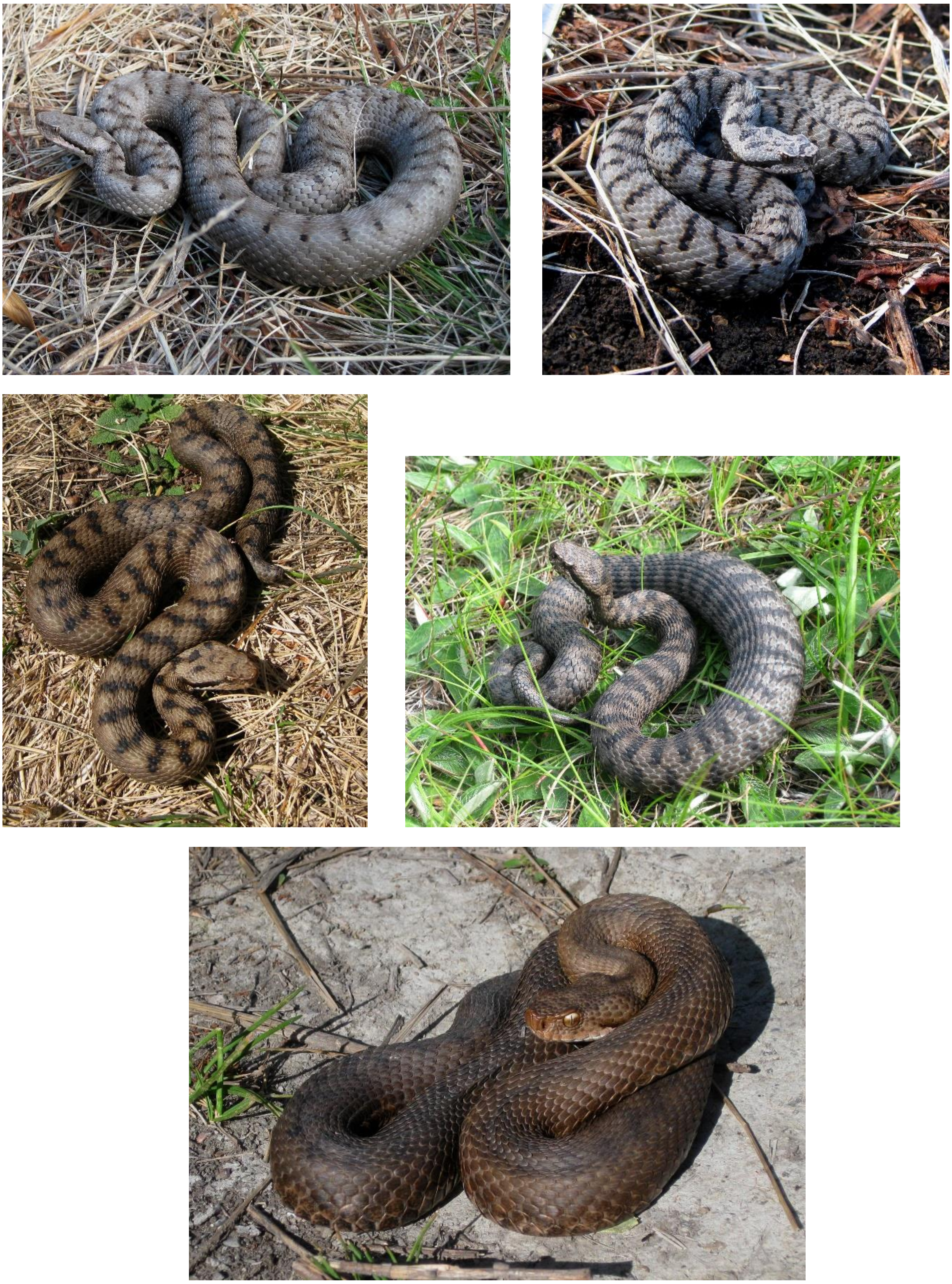

**Figure S3. Progression of accuracy, loss and top3 accuracy through the training period for the three ConvNet of network ensemble: Stochastic Gradient Descent network, Adagrad network and Nadam network.**

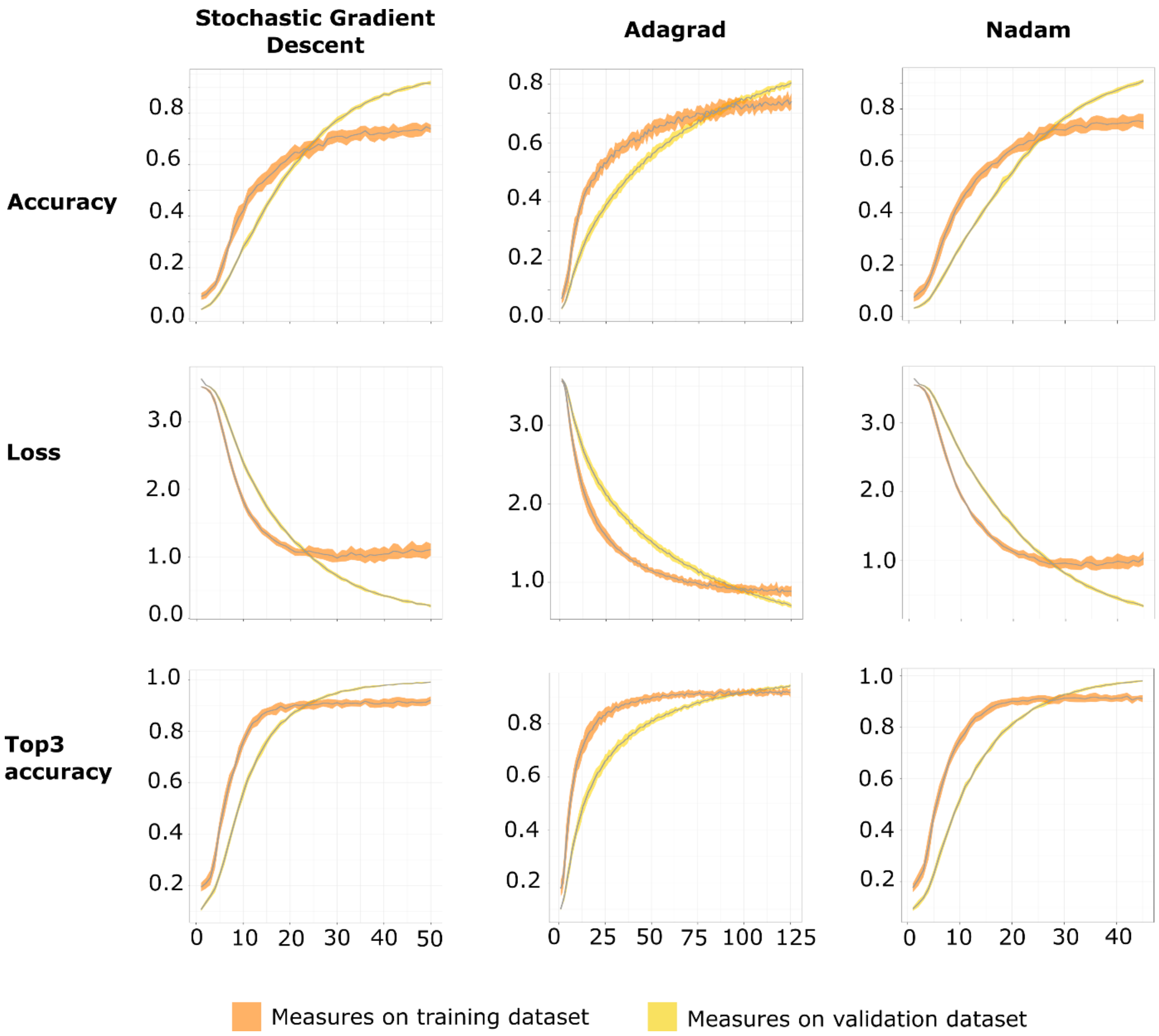
